# PROGMF: Progressive Multimodal Fusion and Representation Learning for Incomplete Neuroimaging Data

**DOI:** 10.64898/2026.11.17.767205

**Authors:** Victor Solomon, Vince D. Calhoun, Tulay Adali, Rogers F. Silva, Rafal A. Angryk, Jingyu Liu

## Abstract

Multimodal neuroimaging studies often involve cohorts with partially overlapping data modalities. Many conventional multimodal fusion methods rely on participants with complete observations across the selected modalities. We introduce Progressive Multimodal Fusion (ProgMF), a novel fusion framework for learning multimodal source representations from incomplete neuroimaging data without imputing missing observations. ProgMF integrates modalities sequentially through independent vector analysis (IVA), cross-stage component alignment, and anchor-guided representation transfer, allowing information learned from earlier-stage cohorts to inform subsequent multimodal decompositions. We evaluated ProgMF on four neuroimaging-inspired synthetic modalities, while varying subject overlap, observation noise, modality relatedness, and fusion orders. ProgMF generally achieved lower source-recovery errors than complete-case IVA and the evaluated imputation-based methods, with larger improvements as multimodal subject overlap decreased. We further applied ProgMF to structural MRI gray matter, resting-state functional network connectivity, and ^18^F-florbetapir (AV45) amyloid positron emission tomography data from the Alzheimer’s Disease Neuroimaging Initiative. ProgMF achieved higher mean cross-modal fusion scores than the imputation-based baselines and a modest improvement over complete-case IVA. Among the 293 participants with complete multimodal observations, two recovered source component vectors showed significant associations with clinical diagnosis across cognitively normal, mild cognitive impairment, and dementia groups. Both sources were also significantly associated with cognitive performance and dementia severity. The recovered sources exhibited diagnosis-related structural and amyloid PET patterns involving regions such as the hippocampi, precuneus, and cingulate cortices. These findings support progressive representation transfer as an effective approach for recovering coherent, interpretable multimodal sources from participants with incomplete modality coverage.

## 1 Introduction

Alzheimer’s disease (AD) is a progressive neurodegenerative disorder that affects memory, cognition, and everyday functioning, ultimately leading to a loss of independence. In 2026, an estimated 7.4 million adults aged 65 years or older in the United States were living with Alzheimer’s dementia, underscoring its substantial public health burden (Alzheimer’s Association, 2026; Scheltens et al., 2021). AD is characterized by the accumulation of amyloid-*β* (A*β*) plaques and tau neurofibrillary tangles, accompanied by synaptic dysfunction and progressive neurodegeneration. However, the onset and progression of these pathological changes vary across individuals (Jack et al., 2024a; Therriault et al., 2024).

This variability has motivated the use of complementary neuroimaging modalities to examine different aspects of AD-related brain alterations. Structural magnetic resonance imaging (sMRI) provides measurements of brain anatomy, including regional gray matter (GM) volume. Resting-state functional MRI (rs-fMRI) measures spontaneous brain activity, from which functional network connectivity (FNC) can be estimated to examine relationships among brain networks. Methods such as NeuroMark have been developed to identify reproducible functional networks from rs-fMRI data (Du et al., 2020; Jack et al., 2024b). Positron emission tomography (PET) using amyloid tracers such as ^18^F-florbetapir (AV45) measures fibrillar A*β* deposition, commonly expressed as standardized uptake value ratios (SUVRs). Integrating GM, FNC, and amyloid PET can therefore provide complementary information about structural, functional, and molecular alterations associated with AD (Ottoy et al., 2025; Landau et al., 2025). Multimodal neuroimaging can also be combined with genetic and epigenetic measurements to investigate biological mechanisms underlying disease-related brain changes (Chen et al., 2026; Duan et al., 2023).

The Alzheimer’s Disease Neuroimaging Initiative (ADNI) provides multimodal neuroimaging, fluid biomarker, genetic, and clinical data from participants across different cognitive and clinical stages, including cognitively normal individuals and those with mild cognitive impairment or dementia (Veitch et al., 2024; Aisen et al., 2024). However, not every participant has every imaging modality acquired at a comparable visit. Differences in acquisition schedules, modality-specific protocols, and access to specialized imaging procedures result in partially overlapping cohorts. Requiring complete observations across multiple modalities can therefore substantially reduce the number of participants available for joint analysis, even when additional observations are available for individual modalities or modality pairs. This limitation is particularly relevant when the objective is to identify reproducible multimodal brain patterns rather than predict a single diagnostic label (Acosta et al., 2022; Hayat et al., 2022).

To address the challenges posed by incomplete multimodal observations, several methods have been developed based on low-rank matrix completion and shared latent representation learning. Matrix completion methods estimate unobserved entries by imposing low-rank structure, often through nuclear-norm regularization (Mazumder et al., 2010). For incomplete multiview clustering, Shao et al. developed a weighted nonnegative matrix factorization approach that accommodates missing views while encouraging agreement among modality-specific latent representations (Shao et al., 2015). Hu and Chen proposed a weighted semi-nonnegative matrix factorization method with basis alignment to learn a common latent representation from incomplete views (Hu and Chen, 2018). Similarly, joint embedding learning and low-rank approximation (JELLA) combines low-rank data approximation with linear transformations to learn shared representations from incomplete multiview data (Tao et al., 2021). In neuroimaging, Liu et al. applied nonnegative matrix factorization to incomplete multimodal ADNI data for participant clustering (Liu et al., 2018). Existing approaches address incomplete multiview data through low-rank reconstruction, shared latent representation learning, or both. Our objective is to estimate corresponding latent sources across modalities while incorporating information from participants who lack complete multimodal observations. To this end, we introduce progressive multimodal fusion (ProgMF), which integrates modalities sequentially and uses source-separation information learned from earlier-stage cohorts to guide subsequent joint decompositions, without imputing missing modality observations.

To implement this progressive fusion strategy, we build upon independent vector analysis (IVA), a blind source separation framework that jointly decomposes multiple datasets by exploiting statistical dependence among corresponding latent sources (Adalıet al., 2014). IVA groups corresponding modality-specific sources into source component vectors (SCVs) and seeks statistical independence between distinct SCVs while allowing dependence among sources within each SCV. Under appropriate conditions on the source distributions and dependence structure, IVA provides identifiability guarantees, including conditions for resolving source permutation ambiguities consistently across datasets (Anderson et al., 2014; Adalıet al., 2014). Recent IVA research has investigated relationships among functional imaging datasets, scalable source estimation, and multimodal representations across dynamic functional states (Lehmann et al., 2024; Gabrielson et al., 2025; Solomon et al., 2026). Building on this foundation, ProgMF independently estimates IVA decompositions at successive fusion stages, aligns corresponding components across stages, and uses previously learned source directions to guide subsequent refinement.

The remainder of the paper is organized as follows. Section 2 presents the proposed ProgMF framework and its optimization procedure. Section 3 describes the datasets, preprocessing methods, evaluation metrics, and experimental setup. Section 4 presents the results from the synthetic data experiments and the ADNI neuroimaging analysis. Section 5 discusses the main findings, methodological implications, limitations, and future research directions. Finally, Section 6 concludes the paper.

## 2 Proposed Method

ProgMF is a multimodal fusion framework designed to learn joint source representations from neuroimaging modalities with partially overlapping subject cohorts. As illustrated in Figure 1, the framework integrates modalities sequentially, beginning with an initial pair and introducing one additional modality at each subsequent stage. At each stage, an IVA model is independently fitted using observations from participants with all currently included modalities. To preserve information learned at preceding stages, corresponding source components are matched through permutation and sign alignment. The aligned unmixing directions are then used as fixed reference anchors in a regularized IVA objective to refine the current-stage source decomposition. Through this process, ProgMF transfers previously learned representations across stages without imputing missing modality observations.

**Figure 1:**
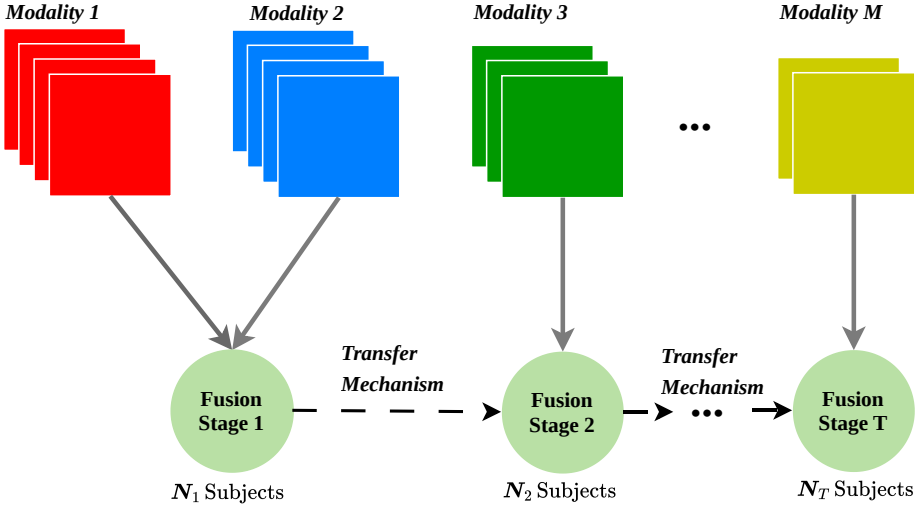
Overview of ProgMF. Modalities are introduced sequentially over nested subject cohorts. Each stage independently estimates a joint source representation, followed by cross-stage component alignment and anchor-guided refinement using directions learned at the preceding stage.

**Figure 2:**
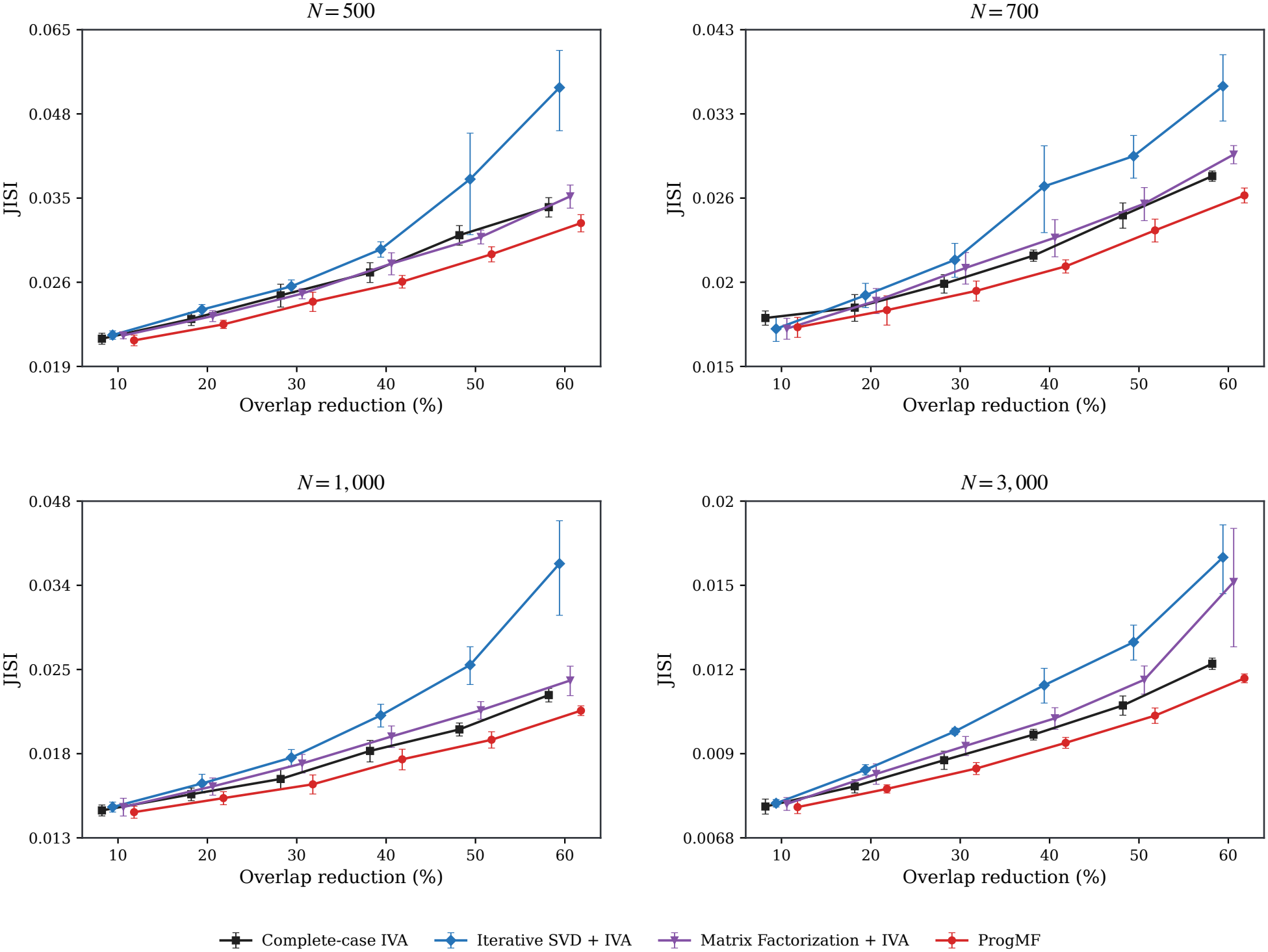
Source recovery under shrinking multimodal overlap. Mean JISI with standard-deviation error bars across ten independent simulation runs for each method and condition. Results are reported at 10 dB SNR for *N* ∈ {500, 700, 1000, 3000}, with overlap reductions ranging from 10% to 60%. The comparison includes complete-case IVA, iterative SVD + IVA, matrix factorization + IVA, and ProgMF.

### 2.1 Problem Formulation

We first formalize the modality availability and subject overlap that determine the successive fusion stages. Consider *M* ≥ 2 imaging modalities, with 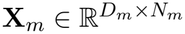 denoting the observation matrix for modality *m*, where *D*_*m*_ is the feature dimension and *N*_*m*_ is the number of available subjects. Let I_*m*_ denote the set of subjects with observations for modality *m*, such that *N*_*m*_ = |I_*m*_|. Because modality availability varies across participants, these subject sets may overlap only partially. Let *ω* = (*m*_1_, …, *m*_*M*_) denote a permutation of the *M* modalities specifying their fusion order. The first stage combines modalities *m*_1_ and *m*_2_, and each subsequent stage introduces one additional modality, resulting in *T* = *M* − 1 fusion stages. At stage *t* ∈ {1, …, *T*}, the active modality set is *M*_*t*_ = {*m*_1_, …, *m*_*t*+1_}, and the corresponding shared cohort is 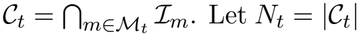 denote the number of subjects with observations from all active modalities. For each *m* ∈ *M*_*t*_, the matrix 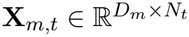 contains the observations for subjects in *C*_*t*_, arranged in a consistent order across modalities. Since *C*_*t*_ ⊆ *C*_*t−*1_ for *t >* 1, the shared cohort size is nonincreasing across fusion stages. The objective is to estimate a joint source representation across all *M* modalities while retaining information learned from the cohorts available at preceding stages. Unlike complete-case fusion, which estimates the entire model using only subjects in *C*_*T*_, ProgMF incorporates earlier-stage information through transferred source-separation parameters rather than reconstructing missing modality observations.

### 2.2 Independent Vector Analysis

ProgMF adopts the transposed formulation of IVA as its underlying linear source-separation model, using the MISA formulatiion (Silva et al., 2021, 2024). In this formulation, each imaging modality is treated as a separate dataset, with imaging features arranged along rows and matched participants along columns. The model estimates modality-specific latent source loadings across participants and captures statistical dependence among corresponding source loadings from different modalities. These relationships are modeled across participants rather than directly between spatial or connectivity patterns. IVA groups corresponding modality-specific sources into source component vectors (SCVs) and seeks statistical independence between distinct SCVs while allowing dependence among the sources within each SCV.

At stage *t* of our fusion framework, each active modality *m* ∈ *M*_*t*_ is associated with an unmixing matrix 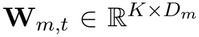, where *K* ≤ *D*_*m*_ is the common number of latent sources per modality, referred to as the model order. The estimated source matrix is given by **Y**_*m,t*_ = **W**_*m,t*_**X**_*m,t*_, where 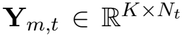 contains the source loadings for the *N*_*t*_ shared participants. Each row of **Y**_*m,t*_ represents the loading profile of one source across participants. Let 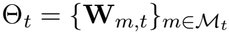 denote the stage-specific model parameters, and let *y*_*m,k,n,t*_ denote the loading of source *k* for participant *n* in modality *m*.

For each source *k*, the corresponding loadings across the *d*_*t*_ = |*M*_*t*_| = *t* + 1 active modalities form an SCV realization for participant *n*, defined as 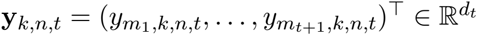. Let **y**_*k,t*_ denote the associated random vector, with the *N*_*t*_ shared participants providing its observed realizations. Each SCV therefore represents a set of corresponding source loadings across modalities, and their joint distribution describes the statistical dependence among these loadings. The IVA objective minimizes the mutual information among the *K* SCVs:

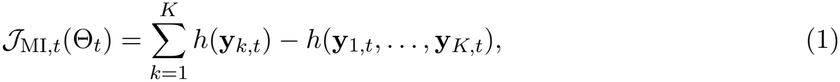

where *h*(·) denotes differential entropy. Minimizing this objective promotes statistical independence between distinct SCVs without imposing independence among the modality-specific sources within each SCV. By exploiting cross-modal source dependence, IVA can establish corresponding source components across datasets under appropriate identifiability conditions (Anderson et al., 2014; Adalıet al., 2014).

To model the joint distribution of the modality-specific sources within each SCV, we adopt the zero-mean multivariate Kotz distribution (Anderson et al., 2013; Silva et al., 2021). The Kotz family provides a flexible distributional model that can capture both second- and higher-order statistical dependence. For SCV *k* at stage *t*, let 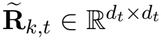 denote a positive-definite dispersion matrix constructed from regularized cross-modal correlation estimates of the current source loadings. The matrix is updated as the source estimates change during optimization. For an SCV realization 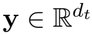, define the quadratic form 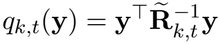, which measures the magnitude of **y** relative to the modeled cross-modal dependence structure.

The corresponding Kotz probability density is

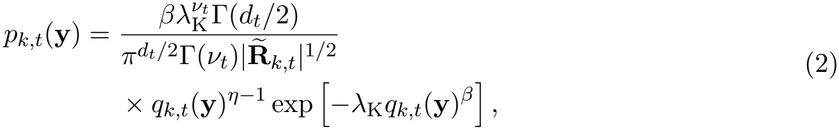

where *v*_*t*_ = (2*η* + *d*_*t*_ − 2)*/*(2*β*), Γ(·) denotes the gamma function, and *β >* 0, *λ*_K_ *>* 0, and *η >* (2 − *d*_*t*_)*/*2 are the Kotz distribution parameters. Specifically, *β* controls the shape of the distribution, *λ*_K_ determines the radial decay, and *η* influences the behavior of the density near the origin. The matrix 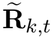 serves as a dispersion parameter describing cross-modal source dependence and is not necessarily equal to the covariance matrix of the resulting Kotz distribution. The notation *λ*_K_ distinguishes the Kotz parameter from the representation-transfer coefficients introduced later.

Using the Kotz density to model the SCV distributions, we obtain a MISA-based source-separation objective comprising the negative log-likelihood of the estimated sources and a correction term for the modality-specific unmixing transformations. For the potentially rectangular unmixing matrices used in ProgMF, we employ the following numerically stabilized objective:

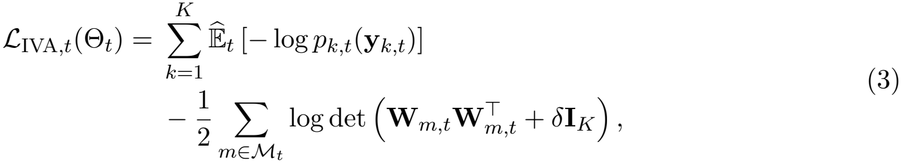

where 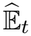 denotes the empirical expectation over the *N*_*t*_ shared participants, **I**_*K*_ is the *K* × *K* identity matrix, and *δ >* 0 is a small numerical stabilization constant. The first term measures how well the estimated SCVs are described by their joint Kotz distributions. The second term accounts for the volume change associated with the unmixing transformations. For full-row-rank rectangular unmixing matrices, the corresponding unstabilized term is equivalent to the sum of the logarithms of their nonzero singular values. The stabilization constant prevents numerical difficulties when the associated Gram matrices approach singularity. Minimizing this objective provides the stage-specific IVA decomposition used by ProgMF.

### 2.3 Progressive Multimodal Representation Learning

Building on the IVA formulation described above, ProgMF estimates joint source representations through successive fusion stages determined by the modality order *ω*. Let FitIVA denote independent optimization of the IVA objective in Equation (3) for a specified set of modalities and source count *K*.

At the first stage, IVA is fitted using the initial modality pair and their shared cohort *C*_1_, yielding 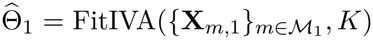). At each subsequent stage *t >* 1, an additional modality *m*_*t*+1_ is introduced, and an IVA model is independently fitted using observations from participants in *C*_*t*_ across all active modalities *M*_*t*_. This produces the initial current-stage estimate FitIVA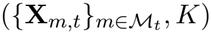.

The preceding estimate 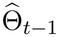 is not used to initialize the current-stage IVA optimization. Instead, independent estimation allows all active modalities, including the newly introduced modality, to contribute to the joint source decomposition before representation transfer. The preceding estimate is subsequently used to guide refinement of 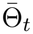. Because 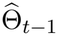 was estimated using the larger or equally sized cohort *C*_*t−*1_, its parameters may retain information from participants who are not included in *C*_*t*_. This information is transferred through previously learned unmixing directions, while the current-stage IVA objective is evaluated using only observations from *C*_*t*_. To ensure that transferred directions correspond to the appropriate current-stage sources, component correspondence is established before refinement.

### 2.4 Cross-Stage Component Alignment

Although IVA jointly establishes correspondence among sources across the active modalities within each stage, independently estimated source decompositions do not necessarily maintain the same component ordering across successive stages. ProgMF therefore establishes cross-stage correspondence using the modalities shared between consecutive stages. This comparison is performed in the common feature space of each inherited modality, with preprocessing transformations maintained consistently across stages. To compare unmixing directions independently of their magnitudes, we define the row-wise normalization operator *N*(**W**)_*i*,:_ = **W**_*i*,:_*/*(∥**W**_*i*,:_∥_2_ + *ϵ*), where *ϵ >* 0 provides numerical stability. For current-stage component *i* and preceding-stage component *j*, their similarity is calculated as 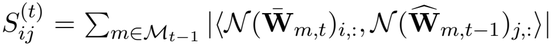, where ⟨·, ·⟩ denotes the Euclidean inner product. This measure aggregates directional similarity across all inherited modalities, allowing corresponding components to be identified using their shared source-separation patterns.

A one-to-one component assignment is obtained by maximizing the total similarity between the current- and preceding-stage decompositions using the Hungarian algorithm. The matched reference directions are then oriented consistently with their current-stage counterparts. We denote the resulting aligned references by 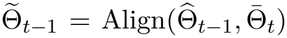, where 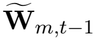 contains the aligned unmixing directions for each inherited modality *m* ∈ *M*_*t−*1_. These references remain fixed during subsequent optimization. This alignment enables representation transfer between independently estimated IVA models without requiring identical subject cohorts or equality of their estimated source loadings.

### 2.5 Anchor-Guided Representation Transfer

Following cross-stage alignment, ProgMF incorporates previously learned source information through an anchor-guided regularization term. The regularizer compares normalized unmixing directions, allowing the current-stage model to adapt to the expanded modality set while encouraging consistency with the preceding representation. For stage *t >* 1, the representation-transfer penalty is defined as

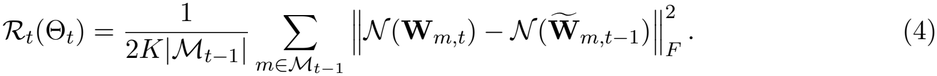

Here, ∥ · ∥_*F*_ denotes the Frobenius norm. The penalty measures the deviation between the current unmixing directions and their aligned references from the preceding stage, averaged across sources and inherited modalities. It applies only to modalities in *M*_*t−*1_ because the newly introduced modality has no preceding reference. Nevertheless, its unmixing matrix is estimated jointly with those of the inherited modalities through the IVA objective. The current-stage optimization combines the IVA source-separation objective with the representation-transfer penalty:

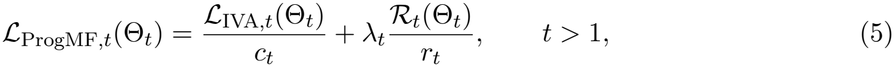

where *λ*_*t*_ *>* 0 controls the strength of representation transfer. The normalization factors are defined as 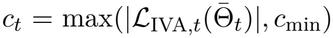 and 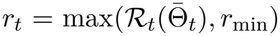, where *c*_min_ *>* 0 and *r*_min_ *>* 0 prevent division by values approaching zero. Both factors are computed using the independently fitted current-stage estimate 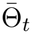 and remain fixed during refinement. The IVA term promotes separation of the joint sources across all active modalities, while the transfer penalty encourages the inherited source directions to remain close to their preceding-stage references. The coefficient *λ*_*t*_ determines the relative influence of these two objectives after normalization. Starting from 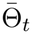, the refined model is obtained as

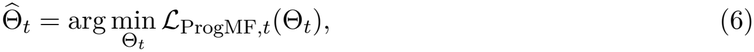

with optimization initialized at the independently fitted parameters. All active modality-specific unmixing matrices are updated during refinement, including the matrix associated with the newly introduced modality. Although the transfer penalty acts only on inherited modalities, the IVA objective models statistical dependence among corresponding sources from all active modalities. Representation transfer therefore provides a soft constraint on the current-stage decomposition rather than fixing previously estimated parameters. This allows ProgMF to incorporate information from earlier cohorts while adapting the joint source representation to additional modalities.

### 2.6 ProgMF Optimization Algorithm

The complete ProgMF procedure is summarized in Algorithm 1. The first stage estimates a joint IVA decomposition from the initial modality pair. Each subsequent stage independently fits IVA using the expanded modality set and its shared cohort, establishes component correspondence with the preceding stage, and performs anchor-guided refinement. The FitIVA operation estimates the stage-specific IVA parameters, Align constructs the corresponding reference directions, and Refine minimizes the regularized objective in Equation (5). The refined parameters are retained as references for the next fusion stage.

#### Algorithm 1

Progressive Multimodal Fusion (ProgMF)

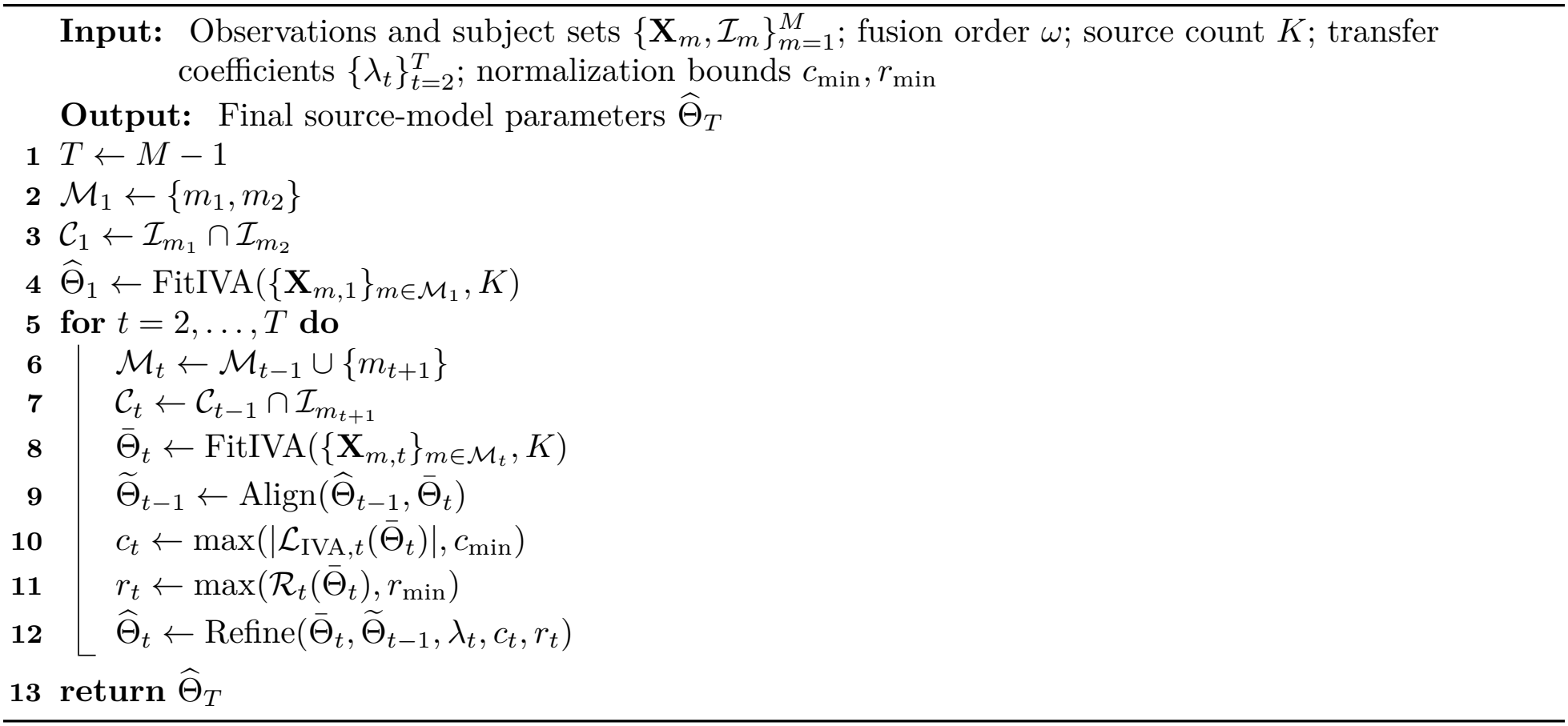

## 3 Materials and Experimental Design

This section describes the synthetic and ADNI neuroimaging datasets, data preprocessing, comparison methods, and experimental procedures used to evaluate ProgMF. The simulation experiments assessed source recovery under varying multimodal subject overlap, observation noise, fusion orders, and early-stage cohort sizes. The ADNI experiments evaluated cross-modal source correspondence and associations of recovered representations with clinical diagnosis, cognitive performance, and dementia severity.

### 3.1 Synthetic Data

Synthetic datasets were generated to evaluate source recovery under controlled variations in sample size, multimodal subject overlap, cross-modal dependence, and observation noise. Four neuroimaging-inspired modalities were considered: structural MRI gray matter (GM), functional network connectivity (FNC), diffusion-derived fractional anisotropy (FA), and positron emission tomography (PET). FNC was represented by the 1,378 unique off-diagonal entries of a symmetric 53 × 53 connectivity matrix, whereas GM, FA, and PET were represented by spatially structured feature vectors. For each modality *m*, the observations were generated using the linear mixing model 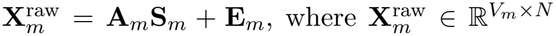 contains observations from *N* subjects across *V*_*m*_ features, 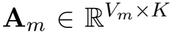 is the modality-specific mixing matrix, **S**_*m*_ ∈ ℝ^*K×N*^ contains the latent sources, and **E**_*m*_ denotes additive Gaussian measurement noise. The simulations used *K* = 10 SCVs. Latent sources were generated using a multivariate Laplace model, allowing statistical dependence among corresponding sources across modalities while maintaining independence between distinct SCVs. For component *k* and subject *n*, the cross-modal source vector was generated as 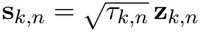, where *τ*_*k,n*_ ∼ Exp(1) and **z**_*k,n*_ ∼ *N*(**0, R**_*k*_). The correlation matrix **R**_*k*_ specified the dependence among the modality-specific sources belonging to component *k*. Modality-specific mixing patterns were constructed using SimTB-derived spatial patterns for GM and PET, smooth tract-like patterns for FA, and structured network connectivity patterns for FNC (Erhardt et al., 2012). Measurement noise was controlled using the signal-to-noise ratio (SNR), defined for modality *m* as 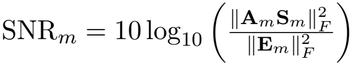, with 10 dB used as the reference condition.

Incomplete modality coverage was simulated using three nested subject cohorts corresponding to successive two-, three-, and four-modality fusion stages. For an initial sample size *N* and cumulative overlap-reduction fraction *r*, the cohort sizes were defined as *N*_1_ = *N*, 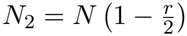, and *N*_3_ = *N* (1 − *r*). Each subsequent cohort was a subset of the preceding cohort, preserving subject correspondence across the available modalities. The primary sample-size experiment considered *N* ∈ {500, 700, 1000, 3000} and overlap reductions up to 60%. The noise-robustness, fusion-order, and early-stage cohort experiments extended overlap reductions to 80%. Complete multimodal observations were generated before restricting modality availability, allowing the ground-truth sources and mixing matrices to be retained for recovery evaluation. Within each simulation realization, all methods were evaluated using the same generated observations and subject-availability patterns. Modality-specific preprocessing transformations were held fixed across fusion stages to maintain consistent feature coordinates for source estimation and cross-stage alignment.

### 3.2 ADNI Neuroimaging Data

Real-data experiments used neuroimaging and clinical data from the Alzheimer’s Disease Neuroimaging Initiative (ADNI), a multicenter study investigating biomarkers and clinical changes associated with Alzheimer’s disease (Veitch et al., 2024). Three imaging modalities were analyzed: T1-weighted structural MRI-derived GM maps, resting-state FNC, and ^18^F-florbetapir (AV45) amyloid PET. Imaging records were matched by participant identifier and acquisition date within a 90-day window. Clinical assessments were linked to the corresponding matched imaging visits. Diagnostic groups were defined using the participants’ current diagnosis at the matched clinical assessment rather than their baseline diagnosis. Clinical measures included the Mini-Mental State Examination (MMSE), the 13-item Alzheimer’s Disease Assessment Scale–Cognitive Subscale (ADAS-Cog13), and the Clinical Dementia Rating Sum of Boxes (CDR-SB). Higher MMSE scores indicate better cognitive performance, whereas higher ADAS-Cog13 and CDR-SB scores indicate greater cognitive impairment and clinical severity, respectively.

Three fusion orders were evaluated: GM + FNC → PET, GM + PET → FNC, and FNC + PET → GM. The corresponding initial two-modality cohorts included 746, 318, and 376 participants, respectively. All three fusion sequences converged to the same 293 participants with matched observations across all modalities. The fully matched cohort comprised 146 cognitively normal (CN) participants, 85 participants with mild cognitive impairment (MCI), and 62 participants with dementia. Demographic and clinical characteristics are summarized in Table 1.

**Table 1:** Demographic and clinical characteristics of the ADNI cohort.

| **Characteristic** | **Overall** | **CN** | **MCI** | **Dementia** |
|---|---|---|---|---|
| Participants, *n* | 293 | 146 | 85 | 62 |
| Age, years | 75.37 ± 7.57 | 74.44 ± 7.81 | 76.06 ± 6.95 | 76.64 ± 7.64 |
| Education, years | 16.39 ± 2.51 | 16.90 ± 2.30 | 16.16 ± 2.73 | 15.48 ± 2.39 |
| MMSE | 27.42 ± 3.37 | 29.11 ± 1.09 | 27.73 ± 2.37 | 23.11 ± 4.17 |
| ADAS-Cog13 | 14.84 ± 11.28 | 8.53 ± 4.25 | 14.20 ± 7.54 | 30.30 ± 11.92 |
| CDR-SB | 1.40 ± 2.22 | 0.13 ± 0.37 | 1.28 ± 1.19 | 4.46 ± 2.75 |

CN, cognitively normal; MCI, mild cognitive impairment; MMSE, Mini-Mental State Examination; ADAS-Cog13, 13-item Alzheimer’s Disease Assessment Scale–Cognitive Subscale; CDR-SB, Clinical Dementia Rating Sum of Boxes.

### 3.3 Neuroimaging Data Preprocessing

#### Structural MRI

T1-weighted structural MRI was represented by voxel-based morphometry GM maps prepared using Statistical Parametric Mapping software (SPM12). Preprocessing included bias-field correction, tissue segmentation, normalization to Montreal Neurological Institute (MNI) space, Jacobian modulation to preserve local tissue volume, and spatial smoothing (Ashburner and Friston, 2005; Sui et al., 2012). Masking was applied to Gray matter images to retain only voxels with intensities ≥ 0.3.

#### Resting-state functional connectivity

Functional network connectivity was derived using the NeuroMark framework implemented in the Group ICA of fMRI Toolbox (GIFT) (Du et al., 2020). The framework estimates subject-specific intrinsic connectivity networks (ICNs) and their corresponding blood-oxygen-level-dependent (BOLD) time courses using predefined spatial templates. We used 53 ICNs distributed across seven functional domains: subcortical, auditory, sensorimotor, visual, cognitive control, default mode, and cerebellar. Following temporal detrending, head-motion regression, and band-pass filtering between 0.01 and 0.15 Hz, static FNC was estimated using Pearson correlations between the ICN time courses. The upper-triangular off-diagonal entries of each symmetric 53 × 53 connectivity matrix were retained, yielding 1,378 features per participant. We refer to this representation as FNC53.

#### Amyloid PET

^18^F-florbetapir (AV45) PET images underwent motion correction, co-registration, frame averaging, spatial standardization, and smoothing (Jagust et al., 2015; Landau et al., 2015). Tracer uptake was expressed as SUVR relative to the cerebellar cortex. The processed SUVR images were restricted using a Harvard–Oxford cortical mask Makris et al. (2006).

#### Dimensionality reduction

Each imaging modality underwent principal component analysis (PCA) and whitening, retaining components that explained at least 85% of the variance. This resulted in 140 retained dimensions for GM, 92 for FNC, and 21 for AV45 PET. The PCA and whitening transformations were estimated using the 293 fully matched participants and projected to the larger cohorts from previous stages of each fusion order.

### 3.4 Model Evaluation

Our evaluation comprised four simulation experiments examining source recovery under varying multimodal overlap, observation noise, fusion orders, and early-stage cohort sizes. Because the ground-truth sources and mixing matrices were known, synthetic recovery accuracy could be evaluated directly. For ADNI, where the underlying sources are unknown, we assessed multimodal correspondence using a correlation-based fusion-quality score. The recovered representations were further examined through associations with clinical diagnosis, cognitive performance, and dementia severity.

#### Simulation evaluation metric

Source-recovery performance was evaluated using joint intersymbol interference (JISI), which measures both the separation of latent sources within each modality and the consistency of corresponding source components across modalities (Anderson et al., 2012). Unlike evaluating source separation independently for each modality, JISI accounts for whether the recovered components share a consistent ordering across datasets. For each modality *m*, the gain matrix is defined as 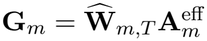, where 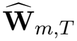 is the estimated final-stage unmixing matrix and 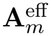 is the ground-truth mixing matrix expressed in the preprocessed feature space. The mean absolute gain matrix is computed elementwise as

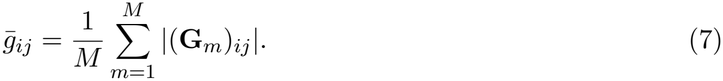

JISI is then defined as

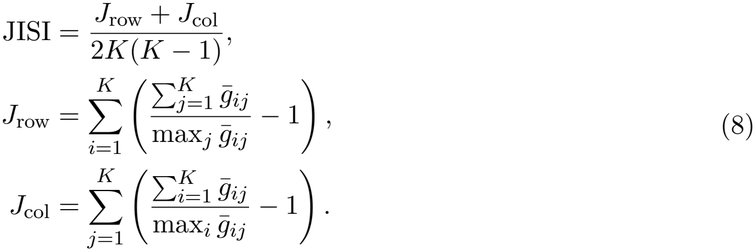

Here, *K* denotes the number of latent sources per modality. The row and column terms quantify residual interference between recovered and ground-truth sources. Because the absolute gain matrices are aggregated across modalities before computing the interference measure, inconsistent source correspondence across modalities also increases JISI. Therefore, a low JISI indicates accurate source separation together with consistent cross-modal component alignment. JISI ranges from zero to one, with zero corresponding to perfect joint source recovery under the assumed mixing model.

#### Neuroimaging evaluation metric

Fusion quality was evaluated by measuring the statistical correspondence between modality-specific sources belonging to the same SCV while accounting for correlations between sources from distinct SCVs. All methods were evaluated using the same 293 participants with complete observations across the three modalities. For *M* = 3, let 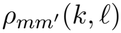 denote the Pearson correlation between source *k* from modality *m* and source *l* from modality *m*^*′*^ across participants. Within-SCV dependence, cross-SCV dependence, and the overall fusion score were defined as

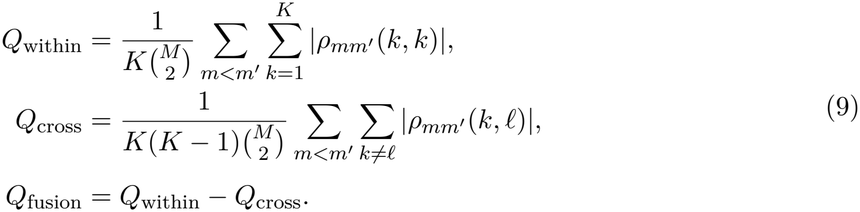

Here, *Q*_within_ measures the average absolute correlation between corresponding sources across modalities, while *Q*_cross_ measures correlations between sources belonging to different SCVs. A higher fusion score indicates stronger within-SCV correspondence relative to cross-SCV correlation. Because the score is based on Pearson correlations, it measures linear source correspondence rather than statistical independence in general.

#### Diagnosis-association analysis

Diagnosis-related variation was evaluated using the three modality-specific source loadings associated with each SCV. A multivariate analysis of covariance (MANCOVA), adjusting for age and sex, was performed for each of the 20 SCVs, using Pillai’s trace as the multivariate test statistic. Bonferroni correction was applied across the 20 joint SCV tests. Individual modality effects were evaluated using age- and sex-adjusted analysis of covariance (ANCOVA), with Bonferroni correction across the 60 modality-specific tests. Pairwise diagnostic contrasts were estimated from the corresponding adjusted models, with correction across the three pairwise comparisons within each panel. These contrasts were treated as follow-up analyses rather than as a separate global family of comparisons.

#### Cognitive and clinical associations

For the selected diagnosis-associated SCVs, associations between modality-specific source loadings and MMSE, ADAS-Cog13, and CDR-SB scores were evaluated using separate linear regression models. Each model was adjusted for age, sex, and education. Source loadings were standardized, and HC3 heteroskedasticity-robust standard errors were used (MacKinnon and White, 1985). Benjamini–Hochberg false discovery rate (FDR) correction was applied across the 18 tested associations.

### 3.5 Comparison Methods

We compared ProgMF with three baseline approaches. Complete-case IVA performs joint source separation using only participants with observations from all modalities.

The iterative singular value decomposition (SVD) + IVA baseline estimates missing observations through repeated low-rank approximations while retaining observed measurements, before applying IVA to the completed data (Troyanskaya et al., 2001). Matrix factorization + IVA approximates the observed data using low-rank factors, reconstructs missing observations, and subsequently applies IVA (Hastie et al., 2015; Mazumder et al., 2010). All approaches used the same IVA source model and model order. Imputation hyperparameters were selected using reconstruction-based criteria without access to ground-truth sources.

### 3.6 Implementation and Experimental Setup

All methods used the same multivariate Kotz source model and a consistent optimization procedure. Model parameters were estimated in double precision using limited-memory BFGS with bounds (L-BFGS-B) (Byrd et al., 1995), with a maximum of 50 iterations per fitting or refinement step. The number of SCVs was fixed at *K* = 10 for synthetic experiments and *K* = 20 for ADNI. ProgMF used strictly positive representation-transfer coefficients. In the primary simulation experiments, these coefficients were selected to minimize JISI for each experimental condition. Because ground-truth sources are available in synthetic data, this constitutes an oracle-guided assessment of source-recovery performance rather than a practical hyperparameter-selection procedure.

The noise-robustness experiments retained the transfer configuration established under the reference condition. For fusion-order sensitivity, the coefficients were fixed at *λ*_2_ = 10^*−*4^ and *λ*_3_ = 10^*−*2^ across all fusion sequences. The ADNI experiments used *λ*_2_ = 10^*−*4^ for the two-stage fusion procedure. The experiments were repeated across ten independent runs per condition to foster robustness. For ADNI, each ProgMF fusion order and baseline method was evaluated across ten independent runs. The GM + PET → FNC configuration achieving the highest fusion score was selected for subsequent diagnosis-association and spatial-pattern analyses. Model selection was based on the unsupervised fusion-quality score without using diagnostic labels. The selected model provided 20 modality-specific source loadings for GM, FNC, and PET across the 293 fully matched participants.

## 4 Results

### 4.1 Source Recovery Under Shrinking Multimodal Overlap

We first evaluated source-recovery performance under decreasing multimodal subject overlap, considering four initial sample sizes, *N* ∈ {500, 700, 1000, 3000}, and overlap reductions ranging from 10% to 60%. The SNR was fixed at 10 dB. Figure 4.1 reports JISI with standard-deviation error bars across ten independent simulation runs for each method and condition.

ProgMF achieved the lowest JISI among the evaluated methods across all reported sample sizes and overlap reductions. Differences between methods were relatively small at higher levels of subject overlap but became more pronounced as the number of fully matched participants decreased. At *N* = 500, ProgMF exhibited a clearer advantage over the baseline methods at 50% and 60% overlap reduction. Similar trends were observed at *N* = 700 and *N* = 1000, with iterative SVD + IVA showing particularly large increases in recovery error. At *N* = 3000, all methods achieved lower recovery errors, although ProgMF continued to yield the lowest mean JISI across the evaluated conditions.

### 4.2 Robustness to Observation Noise

We evaluated source-recovery performance under varying observation noise levels, considering six SNR values, {30, 10, 5, 3, 1, 0} dB, at an initial sample size of *N* = 1000 and overlap reductions ranging from 10% to 80%. Figure 3 reports JISI with standard-deviation error bars for each method and condition. At lower overlap reductions, all methods achieved relatively low recovery errors, with differences becoming more pronounced as the final shared cohort decreased in size.

**Figure 3:**
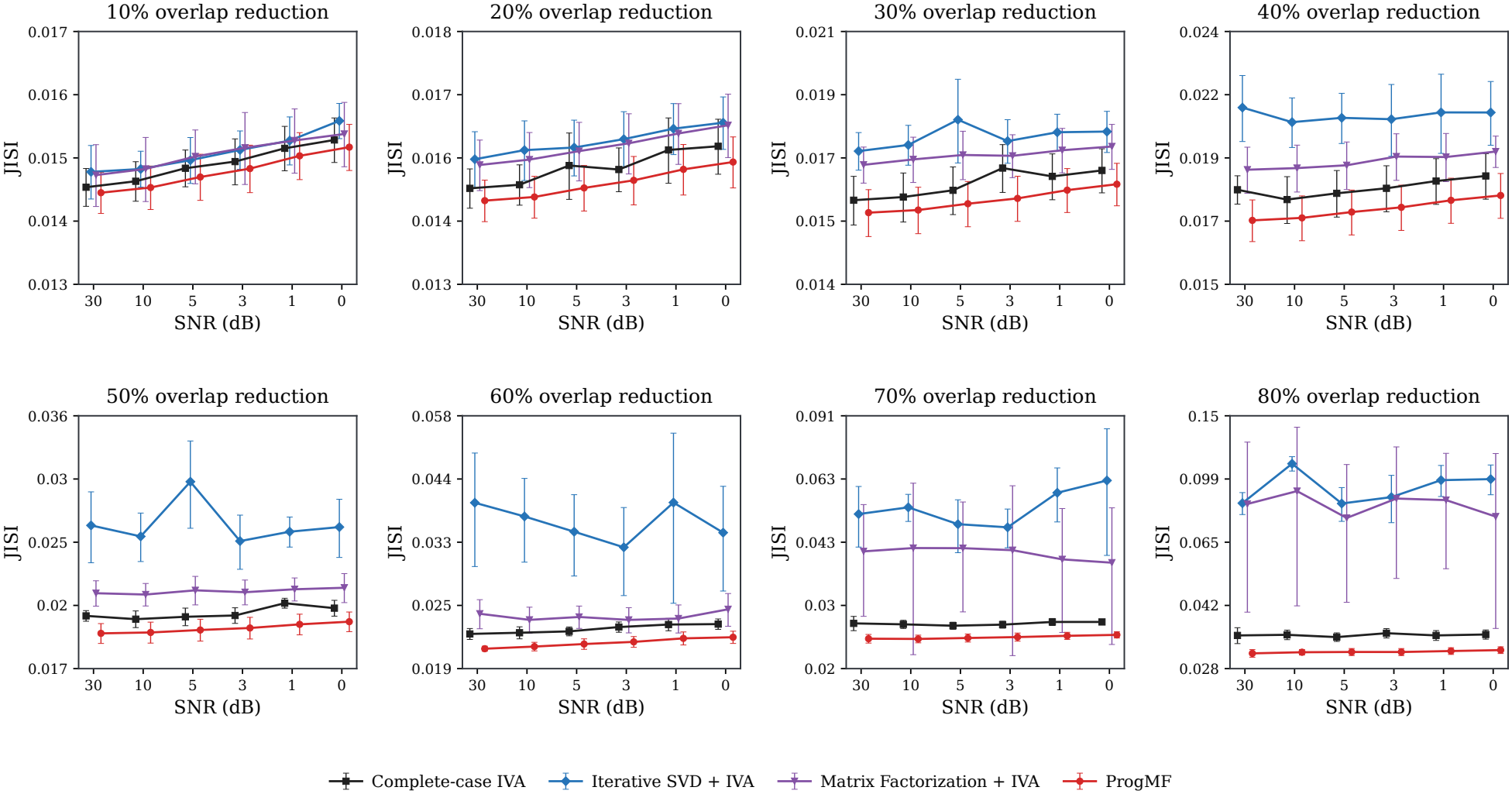
Robustness of source recovery to observation noise. Mean JISI with standard-deviation error bars across ten independent simulation runs for each method and condition. Results are reported for six SNR levels (30, 10, 5, 3, 1, and 0 dB) at *N* = 1000, with overlap reductions ranging from 10% to 80%. The comparison includes complete-case IVA, iterative SVD + IVA, matrix factorization + IVA, and ProgMF.

Across moderate and high overlap reductions, ProgMF generally achieved lower mean JISI and exhibited less variation in recovery performance across SNR levels than the imputation-based baselines. Iterative SVD + IVA showed the greatest variability, particularly under severe overlap reduction, while matrix factorization + IVA also exhibited higher recovery errors. Complete-case IVA remained comparatively stable across noise conditions but generally produced higher JISI than ProgMF.

### 4.3 Sensitivity to Fusion Order

We investigated the effect of modality fusion order on source-recovery performance under heterogeneous cross-modal dependence. The simulation considered strong coupling between GM and FNC, moderate coupling between FA and PET, and weaker dependence among the remaining modality pairs. All 12 distinct fusion orders were evaluated at *N* = 1000 and 10 dB SNR, with overlap reductions ranging from 0% to 80%. Transfer coefficients and optimization settings were held fixed across all fusion orders.

Figure 4 presents the mean JISI with standard-deviation error bars, grouping fusion orders according to the dependence strength of their initial modality pair. Fusion orders beginning with the strongly coupled GM–FNC pair generally achieved lower recovery errors than those initialized with moderately or weakly coupled modalities. At 0% overlap reduction, the strong-coupling group achieved a mean JISI of approximately 0.060, compared with 0.069–0.071 for the moderate- and weak-coupling groups.

**Figure 4:**
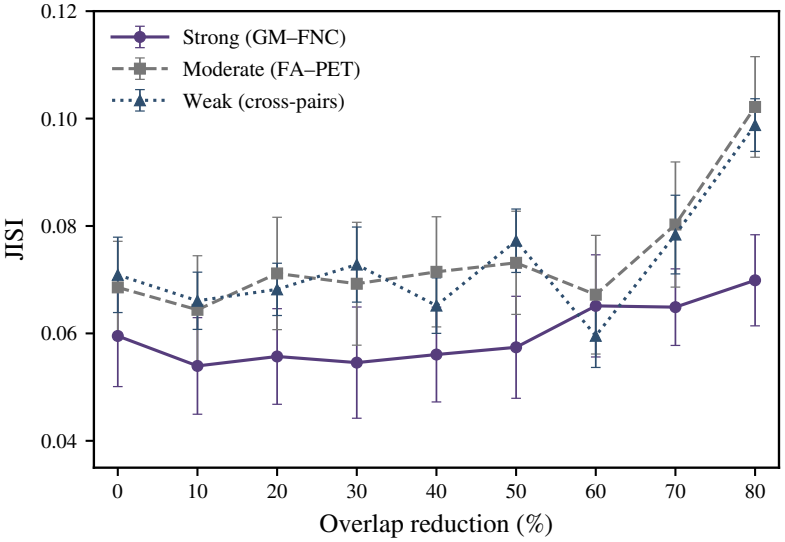
Sensitivity of source recovery to fusion order. Mean JISI with standard-deviation error bars for fusion orders grouped by initial modality-pair dependence: strong (GM–FNC), moderate (FA–PET), and weak (remaining modality pairs). Results are reported at *N* = 1000 and 10 dB SNR, with overlap reductions ranging from 0% to 80%. Transfer coefficients and optimization settings were held fixed across all fusion orders.

### 4.4 Effect of Larger Early-Stage Cohorts

We evaluated the contribution of larger early-stage cohorts by comparing ProgMF with a final-cohort-only progressive variant. Both approaches used the same modality fusion order, representation transfer mechanism, and optimization procedure. Standard ProgMF used the available nested cohort at each fusion stage, whereas the restricted variant used only participants with observations from all four modalities at every stage. This comparison isolates the effect of additional early-stage observations while preserving the progressive fusion procedure.

As shown in Figure 5, ProgMF achieved lower mean JISI across all evaluated overlap reductions. Differences between the two approaches were relatively small at 10%–30% overlap reduction but increased as fewer participants were retained in the restricted early-stage cohorts. At 80% overlap reduction, the final-cohort-only variant yielded a mean JISI of approximately 0.050, compared with 0.022 for ProgMF, representing a 56% reduction in recovery error.

**Figure 5:**
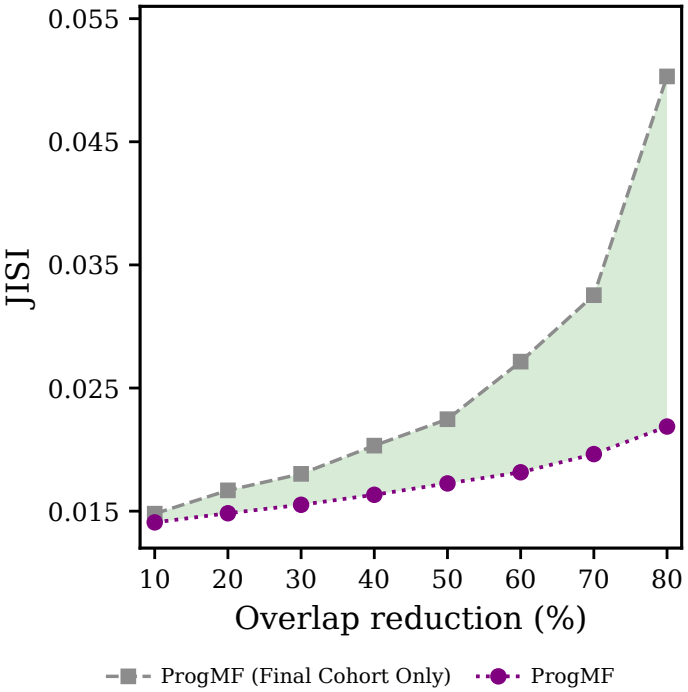
Contribution of larger early-stage cohorts to source recovery. Mean JISI for standard ProgMF and a final-cohort-only progressive variant across overlap reductions ranging from 10% to 80%. Standard ProgMF uses the available nested cohort at each fusion stage, whereas the restricted variant uses only participants with complete four-modality observations throughout all stages. Both approaches use the same fusion order, transfer mechanism, and optimization settings.

### 4.5 Multimodal Fusion Performance on ADNI

We evaluated multimodal fusion performance on ADNI using three fusion orders and three baseline methods. The initial cohort sizes were 746 for GM + FNC, 318 for GM + PET, and 376 for FNC + PET, with all fusion orders converging to the same 293 participantsmwith complete multimodal observations. Table 2 reports *Q*_within_, *Q*_cross_, and *Q*_fusion_ across ten independent runs.

**Table 2:** Multimodal fusion performance on ADNI. Values are reported as mean ± standard deviation across ten independent runs, evaluated on the same 293 participants with complete multimodal observations.

| **Method** | ***Q***<sub>**within**</sub> ↑ | ***Q***<sub>**cross**</sub> ↓ | ***Q***<sub>**fusion**</sub> ↑ |
|---|---|---|---|
| *ProgMF (fusion orders)* |  |  |  |
| GM + FNC → PET | 0.3743 ± 0.0249 | 0.0515 ± 0.0013 | 0.3228 ± 0.0258 |
| GM + PET → FNC | **0.3821 ± 0.0340** | 0.0517 ± 0.0015 | **0.3304 ± 0.0343** |
| FNC + PET → GM | 0.3794 ± 0.0291 | 0.0519 ± 0.0020 | 0.3275 ± 0.0293 |
| Complete-case IVA | 0.3683 ± 0.0198 | 0.0515 ± 0.0011 | 0.3168 ± 0.0207 |
| Iterative SVD + IVA | 0.1076 ± 0.0108 | 0.0508 ± 0.0012 | 0.0568 ± 0.0118 |
| Matrix factorization + IVA | 0.1149 ± 0.0079 | **0.0501 ± 0.0008** | 0.0648 ± 0.0085 |

Among the evaluated fusion orders, GM + PET → FNC achieved the highest *Q*_fusion_ (0.3304 ± 0.0343) and *Q*_within_ (0.3821 ± 0.0340). The other ProgMF configurations achieved comparable *Q*_fusion_ values of 0.3228 ± 0.0258 and 0.3275 ± 0.0293. Complete-case IVA yielded *Q*_fusion_ = 0.3168 ± 0.0207, indicating similar cross-modal correspondence to the progressive configurations. In contrast, iterative SVD + IVA and matrix factorization + IVA achieved substantially lower *Q*_fusion_ values of 0.0568 ± 0.0118 and 0.0648 ± 0.0085, respectively. Values of *Q*_cross_ remained similar across all methods, indicating that the differences in *Q*_fusion_ were primarily associated with *Q*_within_.

### 4.6 Recovered Diagnosis-Related Multimodal Patterns

We examined associations between the ProgMF recovered SCVs and clinical diagnosis in the 293 ADNI participants with complete GM, FNC53, and AV45 PET data. Source-loading associations were evaluated with age and sex included as covariates, and Bonferroni correction was applied across the 60 tested associations. SCV 5 and SCV 2 showed significant associations with diagnosis (Figure 6).

**Figure 6:**
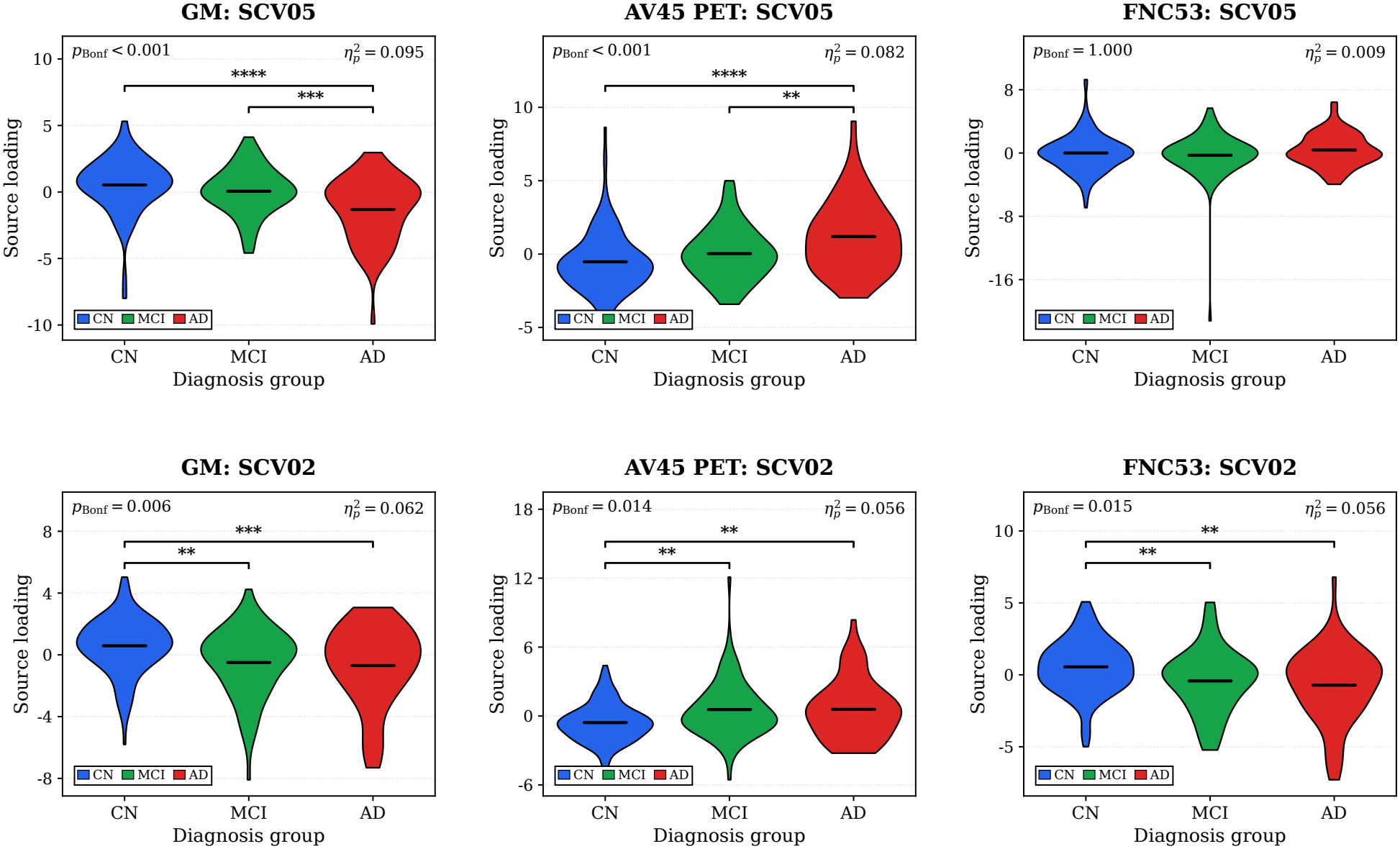
Diagnosis-related source-loading distributions for the recovered multimodal components. Violin plots show source-loading distributions for SCV 5 and SCV 2 across GM, AV45 PET, and FNC53 in 293 participants. Horizontal black lines indicate group means. Significance levels are denoted by ^*^*p <* 0.05, ^**^*p <* 0.01, ^***^*p <* 0.001, and ^****^*p <* 0.0001.

#### Diagnosis-related source-loading associations

SCV 5 showed significant associations with diagnosis in GM (*p*_Bonf_ *<* 0.001, 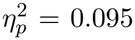) and AV45 PET (*p*_Bonf_ *<* 0.001, *η*^2^ = 0.082). GM source loadings were generally lower in subjects with dementia (AD) than in CN and MCI subjects, whereas AV45 PET loadings showed the opposite pattern. SCV 2 showed significant associations across all three modalities: GM (*p*_Bonf_ = 0.006, 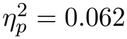), AV45 PET (*p*_Bonf_ = 0.014, *η*^2^ = 0.056), and FNC53 (*p*_Bonf_ = 0.015, 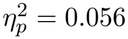). For SCV 2, GM and FNC53 source loadings were generally lower in MCI and AD than in CN subjects, whereas AV45 PET loadings showed the reverse pattern. **Anatomical Patterns and Functional Connectivity of Recovered Multimodal Sources**. We examined the GM and AV45 PET spatial maps of SCV 5 and SCV 2 using the SPM12 Neuromorphometrics atlas (Fonov et al., 2011, 2009) in order to localize how different brain regions contribute to these multimodal patterns our model has recovered. For each map, we thresholded keeping the top 3% of absolute spatial weights. Figure 7 presents ten selected axial slices as representatives of each spatial map.

**Figure 7:**
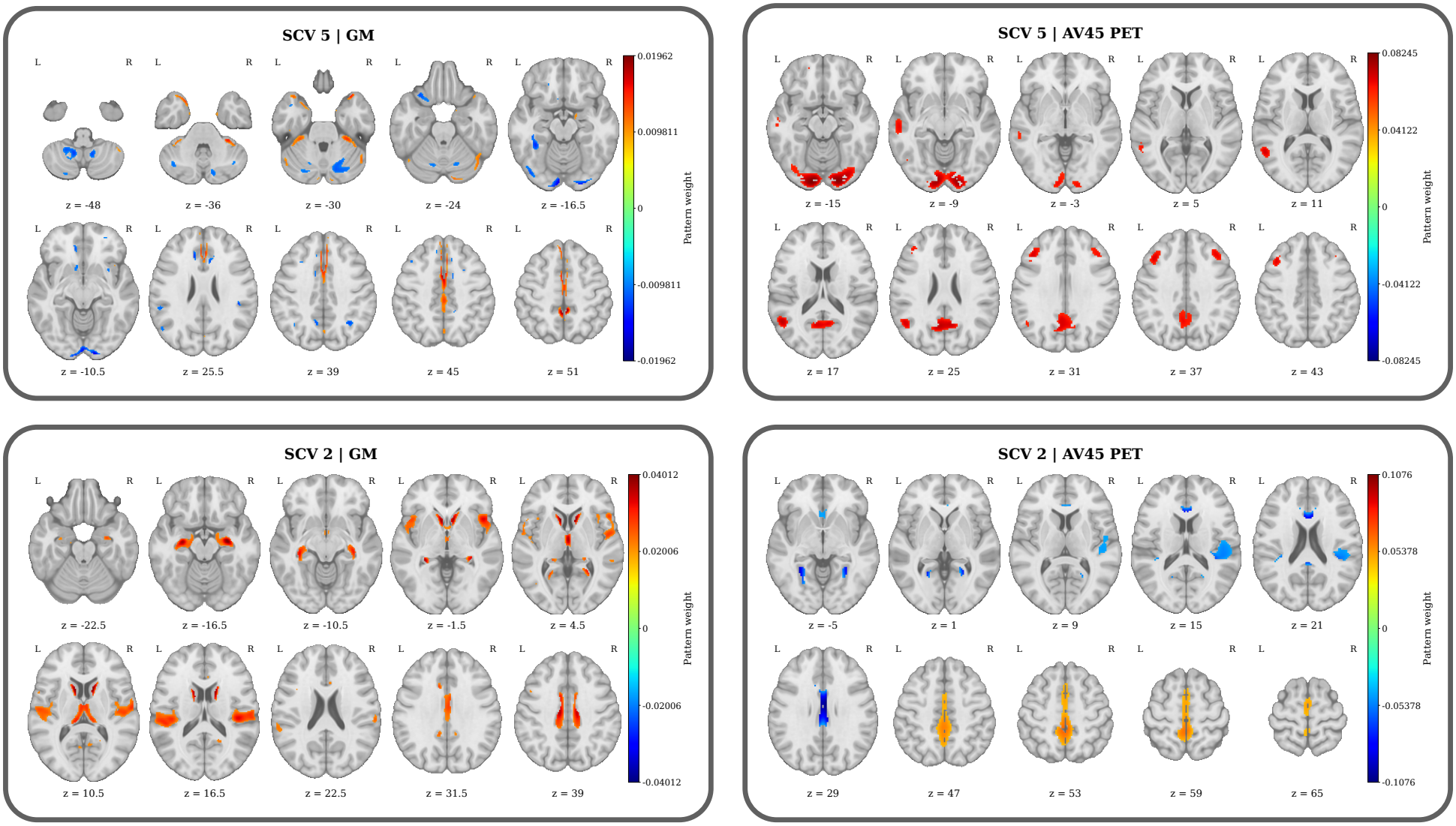
Spatial patterns of the diagnosis-related multimodal components. Selected axial slices from the GM (left column) and AV45 PET (right column) spatial maps for SCV 5 (top row) and SCV 2 (bottom row). The top absolute spatial weights were displayed. Warm colors indicate positive weights, whereas cool colors indicate negative weights.

It can be observed from Figure 7 that SCV 5 shows distributed GM patterns involving cortical and cerebellar regions. Positive weights were observed in cerebellar regions, particularly the right cerebellum, as well as the bilateral supplementary motor cortices, bilateral middle cingulate gyri, left precuneus, and medial segment of the left superior frontal gyrus. These regions contribute to the GM reduction in AD groups. Negative weights were most prominent in the bilateral cerebellum, with additional contributions from the bilateral lingual gyri and left anterior cingulate gyrus. In contrast, all thresholded voxels retained positive weights for AV45 PET in SCV 5, with prominent involvement of the bilateral precuneus, lingual gyri, occipital fusiform gyri, and middle frontal gyri, together with the left middle temporal gyrus. Theses regions correspond to elevated AV45 PET update in AD group.

In the SCV 2 GM map, all top regions retained positive weights, with prominent involvement of the bilateral central opercula, hippocampi, caudate nuclei, and thalami, as well as the left middle cingulate gyrus and left parietal operculum. The SCV 2 AV45 PET map contained both positive and negative weights. Positive weights were concentrated in the bilateral precuneus and supplementary motor cortices, with additional involvement of medial precentral and posterior cingulate regions. Negative weights were most prominent in the right parietal operculum, bilateral middle cingulate gyri, right planum temporale, and right central operculum.

SCV 2 FNC53 showed a distributed pattern of functional connections among the 53 NeuroMark components. Figure 8 presents the 15 connections with the largest absolute weights among the 1,378 possible connections. Prominent positive coefficients included connections between the thalamus and posterior cingulate cortex, cuneus and lingual gyrus, right middle occipital and inferior occipital gyri, and the two hippocampal components. Negative coefficients included connections between the caudate and anterior cingulate cortex, superior and middle frontal gyri, and postcentral gyrus and precuneus. These connections spanned subcortical, visual, cognitive control, default mode, and sensorimotor networks, including both within-network and between-network connections. The coefficients represent contributions to the recovered source, which showed MCI- and AD-related reduction.

**Figure 8:**
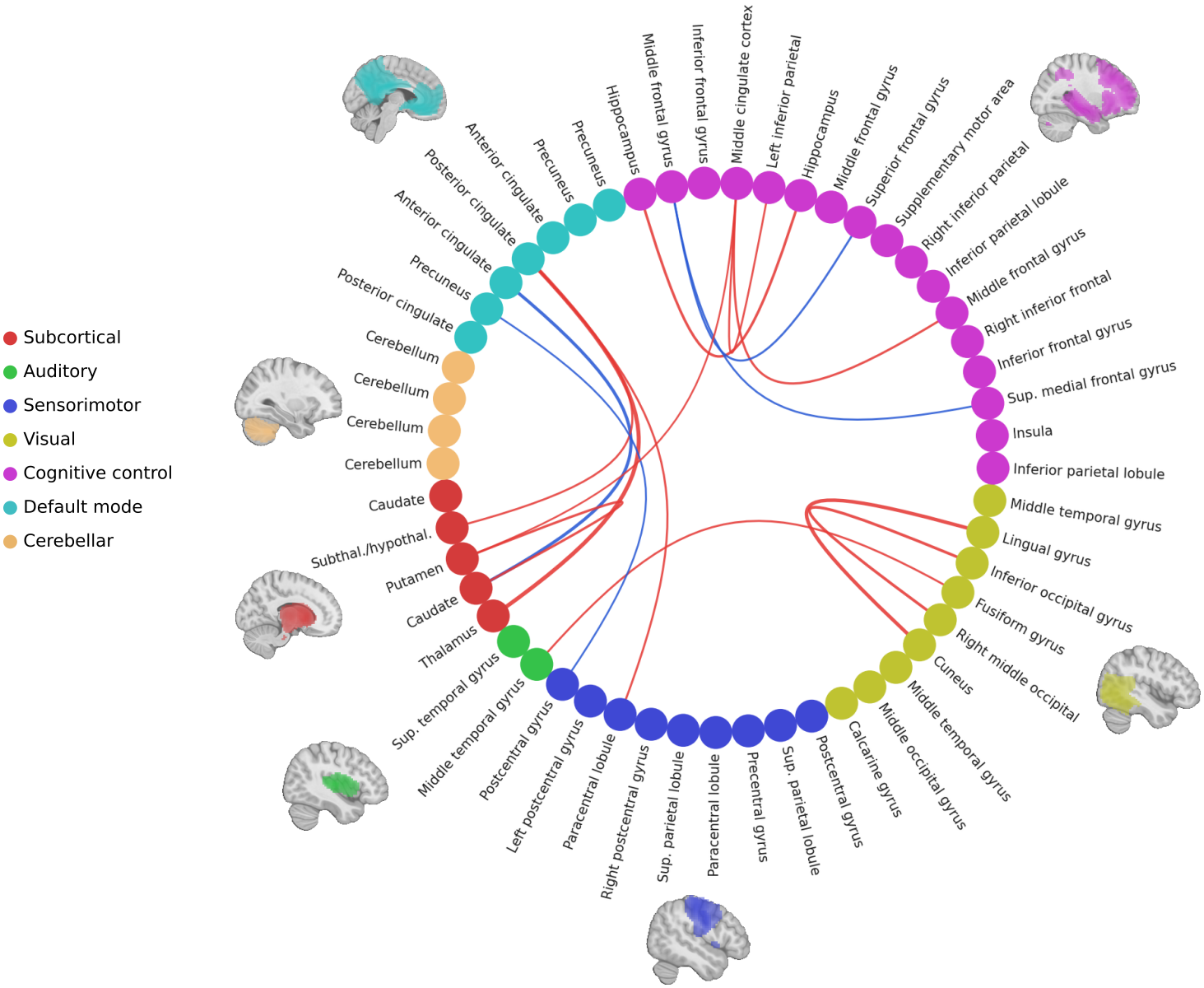
Functional connectivity pattern of SCV 2. Circular connectogram showing the 15 connections with the largest absolute unmixing coefficients in the original FNC53 feature space. The 53 NeuroMark components are grouped into seven functional domains, indicated by the node colors. Red and blue connections represent positive and negative coefficients, respectively, with line thickness proportional to coefficient magnitude.

### 4.7 Associations of Recovered Multimodal Sources with Clinical Cognitive Performance

We evaluated the associations of SCV 2 and SCV 5 source loadings with cognitive performance and clinical severity using MMSE, ADAS-Cog13, and CDR-SB scores in the ADNI cohort. Source loadings were standardized, and HC3 heteroskedasticity-robust standard errors were used MacKinnon and White (1985). Benjamini–Hochberg false discovery rate (FDR) correction was applied across the 18 tested associations.

As shown in Table 3, 16 of the 18 associations remained significant after FDR correction. SCV 2 showed significant associations with all three clinical measures across GM, FNC53, and AV45 PET. Higher GM and FNC53 source loadings were associated with higher MMSE scores and lower ADAS-Cog13 and CDR-SB scores, whereas AV45 PET loadings showed associations in the opposite direction. Similarly, SCV 5 also showed significant associations with all three clinical measures in GM and AV45 PET, with opposite directions between the two modalities. Higher GM loadings were associated with higher MMSE scores and lower ADAS-Cog13 and CDR-SB scores, whereas higher AV45 PET loadings were associated with lower MMSE scores and higher ADAS-Cog13 and CDR-SB

**Table 3:** Associations of SCV 2 and SCV 5 with cognitive performance and clinical severity in ADNI. *β* represents the change in clinical score per standard-deviation increase in source loading, while Std. *β* is the standardized coefficient. scores. For FNC53, only the association with MMSE remained significant after FDR correction.

| **Modality** | **Measure** | *β* | **Std.** *β* | *p* | *q*<sub>FDR</sub> |
|---|---|---|---|---|---|
| **SCV 2** |  |  |  |  |  |
| GM | MMSE | 0.8179 | 0.2425 | 3.2332 × 10<sup>−4</sup> | **5.8197 × 10<sup>−4</sup>** |
|  | ADAS-Cog13 | -3.0883 | -0.2736 | 2.3046 × 10<sup>−5</sup> | **8.2967 × 10<sup>−5</sup>** |
|  | CDR-SB | -0.6586 | -0.2966 | 1.1553 × 10<sup>−3</sup> | **1.7330 × 10<sup>−3</sup>** |
| FNC53 | MMSE | 0.7776 | 0.2306 | 3.0946 × 10<sup>−4</sup> | **5.8197 × 10<sup>−4</sup>** |
|  | ADAS-Cog13 | -2.8984 | -0.2568 | 2.2545 × 10<sup>−4</sup> | **5.0726 × 10<sup>−4</sup>** |
|  | CDR-SB | -0.5558 | -0.2503 | 2.5552 × 10<sup>−3</sup> | **3.2853 × 10<sup>−3</sup>** |
| AV45 PET | MMSE | -0.7750 | -0.2298 | 1.5389 × 10<sup>−3</sup> | **2.1307 × 10<sup>−3</sup>** |
|  | ADAS-Cog13 | 2.9350 | 0.2601 | 1.9310 × 10<sup>−5</sup> | **8.2967 × 10<sup>−5</sup>** |
|  | CDR-SB | 0.6097 | 0.2745 | 4.1031 × 10<sup>−3</sup> | **4.9238 × 10<sup>−3</sup>** |
| **SCV 5** |  |  |  |  |  |
| GM | MMSE | 0.8478 | 0.2514 | 1.3894 × 10<sup>−4</sup> | **3.5728 × 10<sup>−4</sup>** |
|  | ADAS-Cog13 | -2.8771 | -0.2549 | 1.6413 × 10<sup>−5</sup> | **8.2967 × 10<sup>−5</sup>** |
|  | CDR-SB | -0.6671 | -0.3004 | 4.7703 × 10<sup>−6</sup> | **8.2967 × 10<sup>−5</sup>** |
| FNC53 | MMSE | -0.4002 | -0.1187 | 3.9676 × 10<sup>−2</sup> | **4.4635 × 10<sup>−2</sup>** |
|  | ADAS-Cog13 | 0.6621 | 0.0586 | 2.5997 × 10<sup>−1</sup> | 2.7526 × 10<sup>−1</sup> |
|  | CDR-SB | 0.1765 | 0.0794 | 3.0329 × 10<sup>−1</sup> | 3.0329 × 10<sup>−1</sup> |
| AV45 PET | MMSE | -0.6273 | -0.1860 | 1.1226 × 10<sup>−3</sup> | **1.7330 × 10<sup>−3</sup>** |
|  | ADAS-Cog13 | 2.7213 | 0.2411 | 9.3227 × 10<sup>−6</sup> | **8.2967 × 10<sup>−5</sup>** |
|  | CDR-SB | 0.5651 | 0.2544 | 3.6107 × 10<sup>−5</sup> | **1.0832 × 10<sup>−4</sup>** |

## 5 Discussion

This study introduced ProgMF as a progressive approach for learning multimodal source representations from partially overlapping cohorts. The simulation experiments examined source recovery under decreasing subject overlap, observation noise, and different fusion orders, while the ADNI experiments evaluated multimodal correspondence and the clinical relevance of recovered brain patterns. Together, these experiments provide insight into the benefits of transferring previously learned representations and the factors that influence their effectiveness.

### 5.1 Progressive Multimodal Fusion Under Incomplete Data

The simulation results suggest that progressive representation learning can improve source recovery when the number of participants with complete multimodal observations decreases. ProgMF generally achieved lower recovery errors than complete-case IVA and the evaluated imputation-based approaches, with larger differences appearing under more severe reductions in subject overlap. This finding is important for multimodal neuroimaging studies, where requiring every participant to have all imaging modalities can substantially reduce the available sample.

A central feature of ProgMF is that information learned from larger earlier-stage cohorts is retained through aligned source representations rather than reconstructed observations. The comparison with the final-cohort-only progressive variant provides direct evidence for the value of using these additional participants. Both approaches followed the same progressive procedure, but restricting every stage to the final complete cohort resulted in higher recovery errors, particularly at larger overlap reductions. This indicates that the earlier-stage observations contributed useful information to the final decomposition.

The noise-robustness experiments further showed that ProgMF maintained comparatively low recovery errors across the evaluated SNR conditions. Although recovery became more challenging as multimodal overlap decreased, the progressive approach generally remained less sensitive than the imputation-based baselines. These findings suggest that retaining previously estimated source directions can be beneficial when the final multimodal cohort is limited, although the magnitude of this benefit depends on the experimental conditions.

### 5.2 Influence of Fusion Order on Source Recovery

The fusion-order experiments demonstrated that the sequence of modality integration can influence the quality of the recovered sources. In the simulations, fusion orders beginning with the strongly coupled GM–FNC pair generally achieved lower JISI than those initialized with moderately or weakly coupled pairs. One possible explanation is that strongly dependent modalities provide more consistent cross-modal source correspondence at the initial stage, allowing the subsequent transfer procedure to build upon a more reliable representation.

However, the advantage was not uniform across all overlap conditions, indicating that fusion order interacts with the available sample size and dependence structure. Furthermore, the simulated modality pairs differed in both their identities and assigned dependence strengths. Consequently, these results do not establish that stronger initial coupling alone determines the optimal sequence. The ADNI results provide an additional perspective. Although GM + FNC had the largest initial cohort, GM + PET → FNC achieved the highest mean fusion score, with relatively small differences among the three ProgMF orders. This suggests that initial cohort size is not the only consideration when selecting a fusion sequence. The relationships among modalities and the representations learned at successive stages may also influence the final decomposition. Thus, fusion order should be treated as a design choice that warrants evaluation rather than a fixed ordering rule.

### 5.3 Multimodal Representation Quality in ADNI

The ADNI experiments demonstrated that ProgMF could recover corresponding source representations across GM, FNC53, and AV45 PET while incorporating information from partially overlapping cohorts. The progressive fusion orders achieved substantially higher within-SCV correlations than the two imputation-based baselines, whereas cross-SCV correlations remained low across all evaluated methods. These differences suggest that the evaluated imputation-based pipelines were less effective at preserving correspondence between modality-specific sources under the present settings.

Complete-case IVA, however, produced fusion scores relatively close to those obtained with ProgMF. The numerical advantage of progressive fusion was therefore modest in this comparison and does not establish statistical superiority. Nevertheless, ProgMF achieved comparable or slightly higher cross-modal correspondence while making use of observations from participants unavailable in the final complete cohort.

An important consideration is that the proposed fusion-quality metric measures correlation-based source correspondence rather than recovery of known latent sources. Unlike the simulations, ground-truth sources are unavailable in ADNI, and a higher fusion score does not necessarily imply greater biological validity. The real-data results should therefore be interpreted as evidence that ProgMF can maintain coherent cross-modal representations, complementing the source-recovery evaluation performed using synthetic data.

### 5.4 Practical Considerations for PROGMF

The effectiveness of ProgMF depends on the information available at each fusion stage. In applications involving several partially observed modalities, the initial modality pair should be selected by considering both the size of the overlapping cohort and the strength of cross-modal dependence. A larger initial cohort provides more observations for source estimation, while stronger cross-modal relationships may facilitate the recovery of corresponding latent sources. However, neither criterion alone guarantees an optimal fusion order.

We recommend evaluating candidate fusion sequences using consistent model settings and common evaluation subjects. Measures of within-SCV correspondence and cross-SCV separation can help assess the quality of the resulting representations, although these measures do not establish accurate source recovery when ground truth is unavailable. ProgMF is particularly relevant when earlier fusion stages contain substantially more participants than the final fully matched cohort, allowing information from additional observations to contribute through representation transfer. Conversely, when sufficient complete multimodal observations are available, conventional complete-case IVA may provide a simpler alternative without the additional alignment and regularization steps.

### 5.5 Neuroanatomical Relevance of Recovered Multimodal Sources

The spatial patterns recovered by ProgMF provide insight into how structural, functional, and molecular imaging measurements contribute to the same multimodal sources. Although SCV 2 and SCV 5 were both associated with diagnosis, their anatomical distributions differed, suggesting that the learned representations capture distinct aspects of brain variation relevant to Alzheimer’s disease.

SCV 2 GM component exhibited prominent contributions in the bilateral hippocampi, caudate nuclei, thalami, and opercular regions. The significant lower loadings of SCV 2 GM component in AD and MCI groups, compared to controls, indicate that these brain regions all together showed disease related gray matter reduction. Hippocampal involvement is particularly relevant given the established role of medial temporal structures in memory impairment and Alzheimer’s disease (Scheltens et al., 2021; Jack et al., 2024a). The corresponding AV45 PET pattern involved the precuneus, cingulate, and other cortical regions, and these regions contribute to significantly higher loadings in the AD and MCI groups compared to health controls. Our findings are consistent with previous literature suggesting that amyloid-beta accumulation preferentially begins in the precuneus, medial orbitofrontal, and posterior cingulate cortices Palmqvist et al. (2017), before extending to a broader anatomical distribution (Jagust et al., 2015; Landau et al., 2015). The fact that the MCI group also demonstrated significantly increased PET component loadings suggests that amyloid accumulation occurs before the onset of full clinical manifestations of AD, consistent with findings from previous studies (Mormino and Papp, 2018; Jagust and Landau, 2021).

The FNC53 pattern of SCV 2 further involved functional connections among the subcortical, default mode, visual, cognitive control, and sensorimotor networks. Notably, connections involving hippocampal, cingulate, and precuneus appeared as prominent features, while anatomically corresponding regions also contributed to the GM and PET patterns. This spatial alignment suggests that the recovered source captures shared information across anatomically related structural, functional, and molecular measurements.

In contrast, SCV 5 presented a more prominent cerebellar contribution in the GM pattern, together with cingulate, supplementary motor, and precuneus regions. This component showed a more complex pattern of AD-related GM alterations, characterized by lower loadings in regions with positive contributions and higher loadings in regions with negative contributions, primarily in the cerebellum. In comparison, its corresponding AV45 PET pattern revealed a more consistent and spatially distributed pattern of AD-related increased loadings, involving the bilateral precuneus, lingual gyri, occipital fusiform gyri, middle frontal gyri, and temporal gyri. Although SCV 5 involved different brain regions from those identified in SCV 2, both components highlight coherent GM reductions or alterations associated with amyloid-beta accumulation across the continuum from cognitively normal aging to MCI and AD.

### 5.6 Clinical Relevance of Recovered Multimodal Representations

Beyond differences across diagnostic groups, the associations with MMSE, ADAS-Cog13, and CDR-SB provide further insight into the clinical relevance of the representations recovered by ProgMF. Whereas diagnosis-based analyses examine differences among CN, MCI, and dementia groups, these continuous clinical measures capture variation in cognitive performance and disease severity across individuals.

SCV 2 showed consistent associations with all three clinical measures across GM, FNC53, and AV45 PET. The increased GM and FNC53 loadings were associated with better cognitive performance and lower clinical severity in the same direction, whereas the increased PET loadings showed the opposite relationships. This suggests that the different modalities contribute complementary information about clinically relevant variation within the same recovered source. For SCV 5,

GM and AV45 PET loadings, though at different brain regions from SCV 2, were also associated with cognitive performance and clinical severity in a similar way, in addition to the FNC53 weak associations. Together, these findings indicate that clinically relevant variation need not be expressed uniformly across modalities and brain regions. These associations complement the diagnosis-related findings by demonstrating that the recovered representations also relate to continuous measures of cognitive and clinical status.

### 5.7 Limitations and Future Directions

A limitation of ProgMF arises from the assumptions used for cross-stage representation transfer. ProgMF maintains a common number of SCVs across stages and uses one-to-one component matching to align source directions estimated from different subject cohorts. Although this provides a straightforward mechanism for transferring representations, component correspondence may become less reliable when source estimates differ substantially across stages. Furthermore, the current formulation assumes nested subject cohorts, which may not fully accommodate more general patterns of modality availability. Extending ProgMF to support adaptive SCV orders, uncertainty-aware component alignment, and more flexible cohort structures would improve its applicability to heterogeneous multimodal datasets.

The effectiveness of anchor-guided transfer also depends on the regularization coefficients, which control the balance between preserving previously learned representations and adapting to newly introduced modalities. Developing principled data-driven procedures for selecting these coefficients remains an important direction for future research. Finally, ProgMF inherits the linear mixing assumption of independent vector analysis, whereby observed modalities are represented as linear mixtures of latent sources. While this formulation supports explicit source separation and cross-stage alignment, it may not capture nonlinear relationships in complex multimodal data. Future work could investigate nonlinear source-separation models and extend progressive representation transfer to more flexible latent-variable frameworks.

## 6 Conclusion

We introduced ProgMF, a progressive multimodal fusion framework that integrates partially overlapping cohorts through cross-stage alignment and anchor-guided representation transfer without imputing missing modality observations. In controlled simulations, ProgMF generally achieved lower source-recovery errors than complete-case IVA and the evaluated imputation-based approaches, particularly as multimodal subject overlap decreased. The results also demonstrated the contribution of larger early-stage cohorts and the influence of fusion order on source recovery.

When applied to ADNI neuroimaging data, ProgMF recovered corresponding sources across GM, FNC, and AV45 PET, achieving higher mean fusion scores than the imputation-based baselines and modest improvements over complete-case IVA. The recovered multimodal representations revealed GM alterations associated with amyloid accumulation across the continuum from cognitively normal aging to MCI and AD, highlighting their relations with cognitive decline and AD-related clinical manifestation.

## Data and Code Availability

The neuroimaging and clinical data analyzed in this study were obtained from the Alzheimer’s Disease Neuroimaging Initiative (ADNI). ADNI data are available to qualified researchers through the Laboratory of Neuro Imaging (LONI) Image and Data Archive (IDA), subject to approval and the ADNI Data Use Agreement (https://adni.loni.usc.edu/data-samples/adni-data/). Individual participant-level ADNI data cannot be redistributed by the authors.

## Funding

This work was supported by the National Institutes of Health (NIH) under grants R01AG090597 and R01EB006841, and the National Science Foundation (NSF) under grants 2316421 and 2112455.

## Notes

### Competing Interest Statement

The authors have declared no competing interest.

